# Mapping protein neutral networks from predicted secondary structure

**DOI:** 10.64898/2026.03.04.709605

**Authors:** Nabiha Khawar, Sebastian E. Ahnert

**Affiliations:** Department of Chemical Engineering and Biotechnology, University of Cambridge, Cambridge CB3 0AS, UK; The Alan Turing Institute, London NW1 2DB, UK

## Abstract

Protein evolution can be understood as movement through a genotype– phenotype (GP) map, where genetic variation is constrained by the mapping from sequence to structure. While RNA GP maps have been extensively characterised, equivalent maps for proteins remain largely unexplored due to the complexity of folding. Here, we construct an operational GP map for influenza hemagglutinin (HA) using predicted secondary structure as a coarse-grained phenotype.

Combining site-scanning with exhaustive local neighbourhood enumeration, we estimate neutral component (NC) sizes, mutational robustness, and local connectivity patterns across representative HA variants. We observe strong phenotypic bias, with NC sizes spanning orders of magnitude. Robustness increases with estimated NC size, but far more weakly than in RNA, consistent with each NC occupying a vanishing fraction of amino acid sequence space. Local neutral neighbourhood graphs reveal dense, position-centric clusters with limited overlap, suggesting heterogeneous and weakly percolating connectivity. Single-step structural transitions are biased toward structurally similar phenotypes, so that accessible novelty is largely incremental and redundant.

Together, these results provide a tractable empirical framework for protein GP map analysis and suggest that, in HA, neutrality is locally structured yet globally constrained, limiting long-range structural evolvability relative to RNA systems.

## 1 Introduction

Evolutionary outcomes are constrained by the fact that only a subset of phenotypes are both functional and accessible. Viewed through the lens of phenotypic feasibility, certain evolutionary trajectories become statistically more probable than others. The genotype–phenotype (GP) map formalises this bias by defining how genetic variation produces phenotypic outcomes under intrinsic biophysical constraints, independent of environmentally dependent fitness values [19]. Within this framework, sequence space can be represented as a graph in which genotypes are nodes connected by edges linking mutational neighbours that differ by single amino acid substitutions [17].

Each mutation either preserves or alters the phenotype — a binary outcome that forms the conceptual basis of neutral mutations, substitutions that leave structure and thus function unchanged. These neutral mutations form connected networks that permit exploration of sequence space without immediate fitness cost, and their topology determines how populations discover new phenotypes [18, 9]. Studies of RNA sequence–secondary structure maps have revealed general properties of such networks, including redundancy; phenotypic bias, a highly skewed distribution of genotypes per phenotype; neutral correlations, where genotypes of the same phenotype are separated by fewer mutations than expected by chance; and a positive correlation between phenotypic robustness and evolvability [8]. In RNA, large, assortative, and percolating neutral networks make distant phenotypes accessible via sequences of single point mutations [16, 2]. High dimensionality further amplifies these effects, effectively shortening distances between phenotypes and facilitating long-range accessibility [7].

Whether analogous properties exist in real proteins remains unclear. Protein folding involves long-range, context-dependent interactions between chemically diverse residues rather than simple base pairings, which may fragment neutral networks and restrict mutational accessibility. Many sites in proteins are intolerant to substitution due to their functional specificity, producing locally rugged fitness landscapes [15]. Compensatory epistasis can sometimes alleviate these constraints, but the order of substitutions and the connectivity of neutral networks then become decisive determinants of which evolutionary paths remain accessible [13, 3].

Until recently, limited predictive accuracy restricted such studies in proteins to small systems or coarse-grained models such as lattice proteins. Advances in co-evolutionary inference [12, 14] and deep learning–based structure prediction [21, 20] now enable large-scale, quantitative exploration of protein sequence–structure relationships. Using predicted secondary structures as phenotypic outcomes provides a computationally tractable representation that captures key aspects of conformational variability. Although coarse-grained, this physically grounded description uses structural states that constrain tertiary organisation while abstracting away finer details such as side-chain packing.

Here, we develop a conceptual and empirical extension of GP map theory for a real protein system: influenza hemagglutinin (HA), a highly variable yet structurally constrained protein. Using predicted secondary structure to impose biophysical constraints, we statistically infer robustness, connectivity, and evolvability from local sequence samples, approximating how structural constraints organise accessible sequence space.

## 2 Methods

### 2.1 Protein sequence data

HA sequences were obtained from public influenza databases (NCBI, GISAID). Only full-length sequences *L* = 566 without ambiguous residues (“X”) were retained, resulting in 19,289 sequences. Each residue was assigned one of three secondary structure states (C = coil, H = *α*-helix, E = *β*-sheet) using Porter5 [21]. Despite the large number of input sequences, only 2,429 unique secondary structure strings were observed. Predicted secondary structures were validated against experimental PDB structures for three representative strains, showing high concordance (Q3 ≈ 0.74–0.77, See Supplementary Table S1).

To reduce temporal sampling bias in the dataset towards recent isolates, we retained the five most frequent secondary structures per year, yielding 197 representative phenotypes. From this set, we selected fifteen phenotypes that covered the full observed range of neutral mutant counts identified through global exploration for detailed analysis.

### 2.2 Definition of the GP map

Let 𝒢 denote the full amino acid sequence space of HA (|𝒢| = 20^566^). The GP map is defined as *f* : 𝒢 → Φ, which assigns to each sequence *g* ∈ 𝒢 a secondary structure string *ϕ* = *f* (*g*) ∈ ⊕. The neutral set of a phenotype *ϕ* is the set of sequences mapping to *ϕ*, and an NC is a maximally connected subset of this set under single-point mutations. Since HA has length *L* = 566, exact characterisation of its NCs is computationally infeasible; instead, we estimate their properties using statistical methods adapted from previous studies [6, 11, 5, 24].

We focus on three descriptors: **Size**, the estimated number of genotypes realising a phenotype; **Robustness**, the fraction of single site mutations that preserve the phenotype [7]; and **Topology**, the internal connectivity of genotypes within an NC [2].

### 2.3 Estimation of NC size and robustness

NC size and robustness were estimated using site versatility, which measures the effective number of amino acids tolerated at each position without altering the phenotype. Let 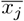 denote the mean number of neutral substitutions observed at site *j* across the sampled genotypes. The NC size estimator is then given by

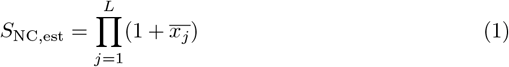

where the +1 term accounts for the resident amino acid at each site. This calculation assumes approximate independence of mutational tolerance across sites, ignoring higher-order epistatic interactions. Given that epistasis is pervasive in proteins, *S*_NC,est_ should be interpreted as an effective combinatorial approximation, with relative differences and scaling relationships more robust than absolute values.

Robustness, the probability that a random point mutation is neutral, was estimated as

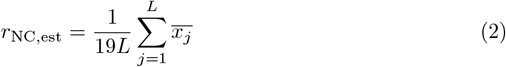

The site versatility values that feed these estimators were obtained by combining global and local exploration strategies.

For global exploration, we implemented a deterministic site-scanning procedure. Starting from a reference genotype, sites were scanned sequentially (0–*L* − 1). At each site, the 19 alternative amino acids were tested in randomised order until a neutral substitution was identified. The secondary structure of each mutant was predicted and compared to that of the parent; only mutants maintaining identical predicted structure were retained. Each newly accepted neutral genotype then served as the starting point for the next site, allowing progressive expansion of the explored subset of the NC. After the final site, scanning resumed from site 0 using the most recently accepted neutral genotype. At least three full iterations were performed. Since neutrality was defined by strict structural identity, prediction errors could only reduce measured robustness and increase apparent fragmentation of the NC.

For local exploration, we exhaustively enumerated all 19 alternative amino acids at each site for a given genotype, generating 19*L* = 10,754 single-point mutants per sequence. This procedure was applied to random subsamples of genotypes of size *S* ∈ {10, 20, 50, 100}, with *R* = 2 independent replicates. NC size estimates converged rapidly with increasing *S* (coefficient of variation ≈ 0.015 at *S* = 10), and *S* = 20 was therefore adopted for all subsequent analyses (see Supplementary Figure S1).

Specifically, NC size estimation uses exhaustive enumeration of single-point neighbourhoods for a random subset of 20 genotypes sampled from the site-scanning data, following the approach validated by Weiß and Ahnert [24] for RNA secondary structure.

### 2.4 Coarse NC approximation directly from site-scanning

As a faster alternative to exhaustive enumeration, per-site versatilities can be estimated directly from site scanning by recording the number of attempts required to find a neutral mutant at each site.

For site *j*, let *A*_*ij*_ denote the index of the first neutral substitution in a randomised ordering of the 19 alternative amino acids during scanning loop *i*. Under sampling without replacement, *A*_*ij*_ follows a negative hypergeometric distribution with expectation

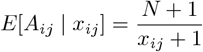

where *x*_*ij*_ is the number of neutral amino acids at site *j* and *N* = 19.

A method-of-moments estimator for *x*_*ij*_ is therefore

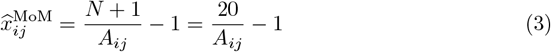

Sites with no observed neutral substitutions (i.e. *A*_*ij*_ = *N*) were assigned a small pseudocount (*x*_*ij*_ = 0.05). Since scanning terminates at the first neutral substitution, a single loop underestimates versatility; repeated loops canalise exploration toward more connected regions of the NC, improving the estimates. The mean site-specific versatility across *i* site scanning loops was calculated as

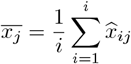

Per-site robustness was then estimated as

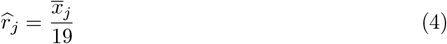

which is bounded between 0 and 1.

Robustness estimates from site scanning exhibit bimodal distributions reflecting extreme heterogeneous constraints across sites (Figure 1a). However, because scanning stops at the first neutral mutation, early successes can yield robustness values of 1, which are clear overestimates. By contrast, exhaustive enumeration evaluates all substitutions and provides more accurate local tolerance estimates, though it is computationally feasible only for smaller samples. Consequently, these estimates reflect local properties of the NC and may vary across sampled genotypes, particularly in highly heterogeneous components (Figure 3b). Divergences between the methods highlight the scale-dependent structure of protein neutral networks, where local and global neutrality need not coincide due to variation in network density.

**Figure 1.**
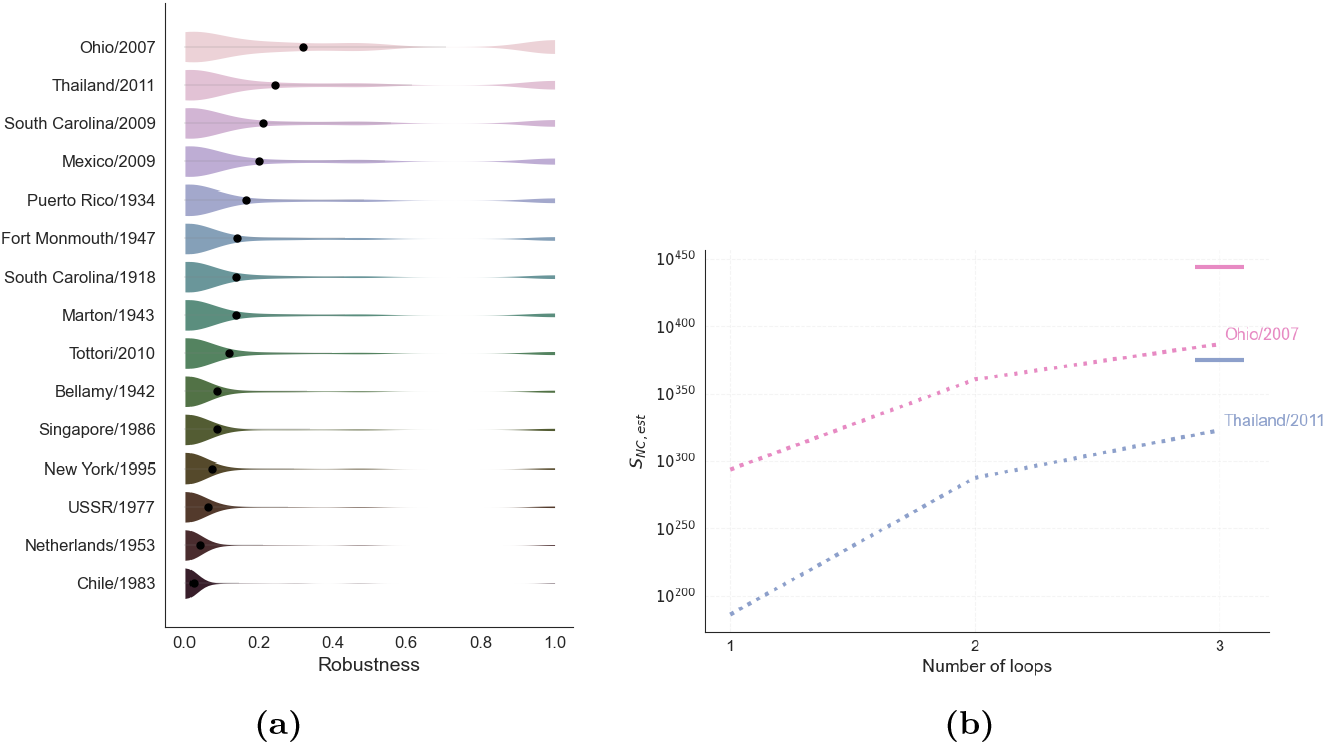
Site-scanning based estimations: (**a**) Robustness density for each strain estimated via site-scanning. (**b**) NC size estimates from site scanning (dashed lines) using 1, 2, and 3 scanning loops, compared with the estimate from exhaustive neighbourhood enumeration (horizontal line). Increasing the number of loops improves agreement for both Ohio/2007 and Thailand/2011 strains.

While exhaustive enumeration provides stable NC size estimates, site-scanning offers a faster approximation. Single-loop estimates tend to underestimate NC size, but increasing the number of loops progressively improves agreement with exhaustive enumeration (Figure 1b).

### 2.5 Network reconstruction and topology analysis

To characterise the local structure of NCs, we reconstructed partial NC graphs from sequences obtained by exhaustively enumerating the one-step neighbourhoods of 20 seed genotypes per strain. For each strain, all seed genotypes and all their neutral single-point mutants were pooled to define the node set of a strain-specific graph.

Edges were placed between any pair of sequences that differed by a single amino acid substitution, yielding an undirected graph representing local neutral mutational connectivity. Although these graphs capture only a subset of the full NC, they preserve the immediate connectivity relationships among sampled genotypes, enabling estimation of local topological properties.

Network properties, including degree distributions and component structure, were analysed using the NetworkX Python library.

## 3 Results

Using exhaustive single-step neighbourhood enumeration of 20 randomly subsampled genotypes per strain (Section 2.3), we quantified NC size and robustness, revealing that larger networks are marginally more mutationally tolerant. We then reconstructed local NC topology (Methods 2.5) to examine connectivity patterns, showing that networks are generally star-like and modular, with highly connected seed genotypes bridging weakly overlapping neighbourhoods. Finally, we assessed evolvability by enumerating all non-neutral single-point substitutions for these genotypes, finding that structurally accessible phenotypes are highly redundant and clustered near the reference structure, highlighting that HA’s capacity for phenotypic innovation is both local and constrained.

### 3.1 Estimated NC sizes and global robustness

Across strains, *S*_NC,est_ spans more than two hundred orders of magnitude, 10^176^–10^444^, indicating a strong phenotypic bias in HA structure space. A small number of secondary structures dominate sequence space, while most occupy comparatively small regions, even among those observed in consensus sequences. Ranking NCs by decreasing size produces a highly skewed, Zipf-like rank–size relationship (Figure 2a). As observed for RNA secondary structures by Schuster et al. [17], this approximate scaling is confined to a restricted range of ranks and deviates from scale-free expectations. Specifically, the distribution flattens at intermediate ranks and decays rapidly in the tail.

**Figure 2.**
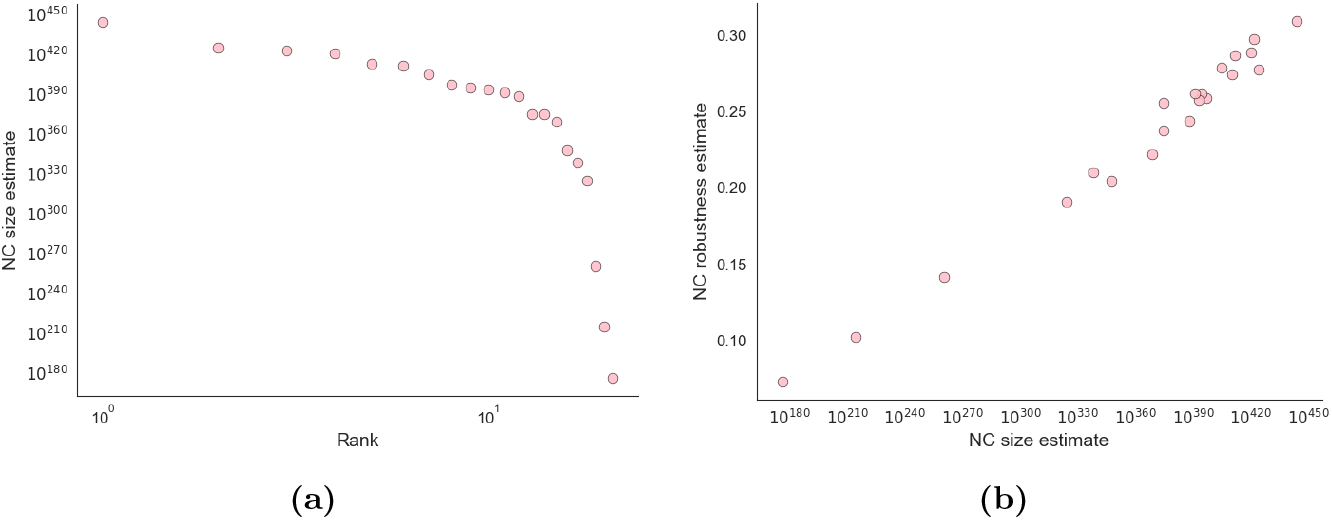
NC properties computed using exhaustive neighbourhood enumeration: (**a**) Rank–size distribution of HA protein NCs obtained from exhaustive single-step neighbourhood enumeration. (**b**) Robustness–size relationship of HA protein NCs obtained across all strains.

Larger NCs were also more mutationally robust (Figure 2b), consistent with the canonical scaling observed in RNA GP maps [10, 2, 8]. In HA, robustness increased monotonically with size (Pearson’s *r* = 0.991, *p* = 4.4 × 10^−18^; Spearman’s *r* = 0.973, *p* = 1.6 × 10^−13^), but the slope of the relationship (*β* = 0.001) was markedly shallower than that reported for RNA (*β* ≈ 0.1, [10]). This difference can be derived analytically: robustness scales with the fraction of genotype space occupied by a neutral network, *S*_NC_*/k*^*L*^ which depends sensitively on alphabet size *k* and sequence length *L*. Since HA occupies an infinitesimal portion of its vast genotype space (|𝒢| = 20^566^), its neutral networks are sparse and boundary-dominated, yielding a greatly reduced *β*. Thus, although strains with larger neutral networks are somewhat more mutation-tolerant, the gain in robustness is minimal, reflecting the inherent rigidity of protein structure space.

To examine how robustness varies across strains, we plotted the number of attempts required to identify a neutral mutant against cumulative site-scanning attempts, using this as a measure of local mutational constraint. Discovery curves differed markedly between strains. Highly constrained NCs, such as *Netherlands/1953*, produced flatter discovery curves, while less constrained NCs, including (*Mexico/2009, Ohio/2007*), exhibited smoother, more continuous increases (Figure 3a).

**Figure 3.**
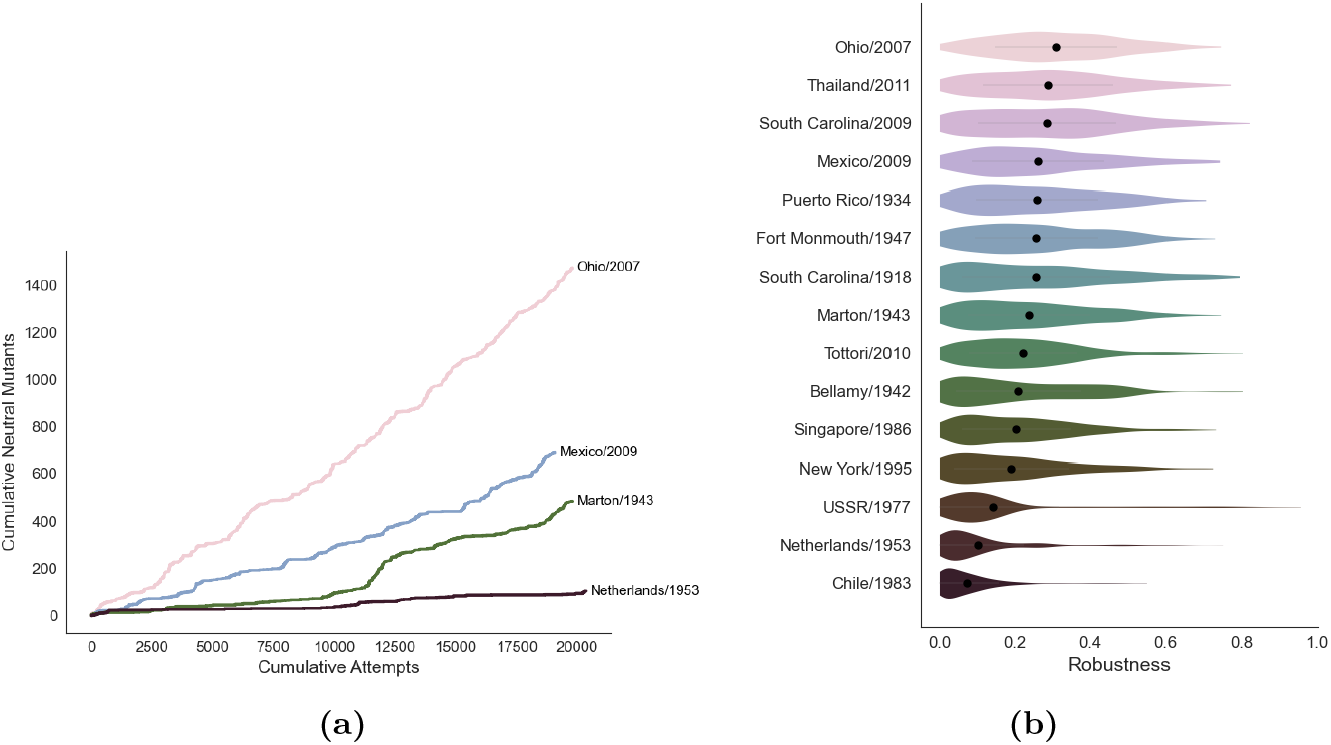
Strain-dependent variation: (**a**) Cumulative number of neutral mutants found versus mutation attempts during site scanning for four representative phenotypes. **(b)** Robustness density for each strain obtained via exhaustive neighbourhood enumeration.

Absolute site-specific robustness values, obtained via exhaustive enumeration of the 20 genotypes per strain (Figure 3b), also revealed strain-dependent variation. Some strains exhibited multimodal distributions, with a high density of sites occupying distinct robustness values, whereas others were more uniform, with only a thin tail extending toward higher robustness.

Such variation in strain-specific robustness suggests that neutral mutations are unevenly distributed within sequence space. Some NCs are densely connected, allowing frequent neutral exploration, whereas others are sparse and fragmented, constraining the paths populations can traverse. To characterise these patterns more explicitly, we next examined the local topology of NCs.

### 3.2 Local topology of NCs

Table 2 summarises the properties of five representative NCs. All display a characteristic star-like topology, in which neighbour nodes have low degrees and connect primarily to their seed genotype or to variants differing at the same position. In all cases, the number of connected components is smaller than the number of sampled seed neighbourhoods, indicating partial overlap between neighbourhoods.

**Table 1.** Estimated NC sizes and robustness for selected HA strains using exhaustive local neighbourhood enumeration of 20 sequences using Equation 1 and 2 respectively.

| Strain | Size ($x, 10^x$) | Robustness |
| --- | --- | --- |
| Ohio/2007 | 443.77 | 0.309 |
| Mexico/2009 | 420.07 | 0.289 |
| Tottori/2010 | 411.88 | 0.287 |
| South Carolina/1918 | 396.79 | 0.259 |
| Fort Monmouth/1947 | 394.39 | 0.262 |
| Puerto Rico/1934 | 392.99 | 0.257 |
| Marton/1943 | 374.65 | 0.256 |
| Thailand/2011 | 374.63 | 0.238 |
| South Carolina/2009 | 368.62 | 0.222 |
| Bellamy/1942 | 347.31 | 0.204 |
| USSR/1977 | 337.95 | 0.210 |
| New York/1995 | 324.35 | 0.191 |
| Singapore/1986 | 260.52 | 0.142 |
| Netherlands/1953 | 214.54 | 0.103 |
| Chile/1983 | 176.26 | 0.073 |

**Table 2.** Properties of five representative reconstructed NCs, showing total node counts, number of components, and largest component size.

| Strain | Nodes | Comps. | Largest (%) |
| --- | --- | --- | --- |
| Mexico/2009 | 62 041 | 17 | 11.8 |
| South Carolina/1918 | 55 636 | 16 | 13.9 |
| USSR/1977 | 45 140 | 12 | 40.4 |
| New York/1995 | 41 123 | 15 | 17.8 |
| Netherlands/1953 | 22 061 | 11 | 27.9 |

Since the neighbourhoods of the neighbours were not exhaustively enumerated, these graphs necessarily underestimate long-range connectivity and appear more fragmented than the full NCs. Despite this limitation, relative differences in local topology remain informative, as neighbourhood overlap and local degree structure have been shown to predict global connectivity in other GP maps.

Substantial variation in local connectivity is observed across strains. The fraction of nodes contained within the largest connected component ranges from below 15% in several strains to over 40% in *USSR/1977*. Larger NCs do not necessarily exhibit more connected dominant components, and local fragmentation appears to be a general property of HA NCs.

A small number of nodes exhibit extremely high degree, corresponding to the seed genotypes from which neighbourhoods were exhaustively enumerated. Degree distributions restricted to seed genotypes (Figure 4a) show that smaller NCs tend to have narrower neighbourhoods, reflecting reduced positional tolerance. *USSR/1977*, in particular, exhibits substantial heterogeneity across its 20 seeds, indicating marked variation in local robustness even within a single NC.

**Figure 4.**
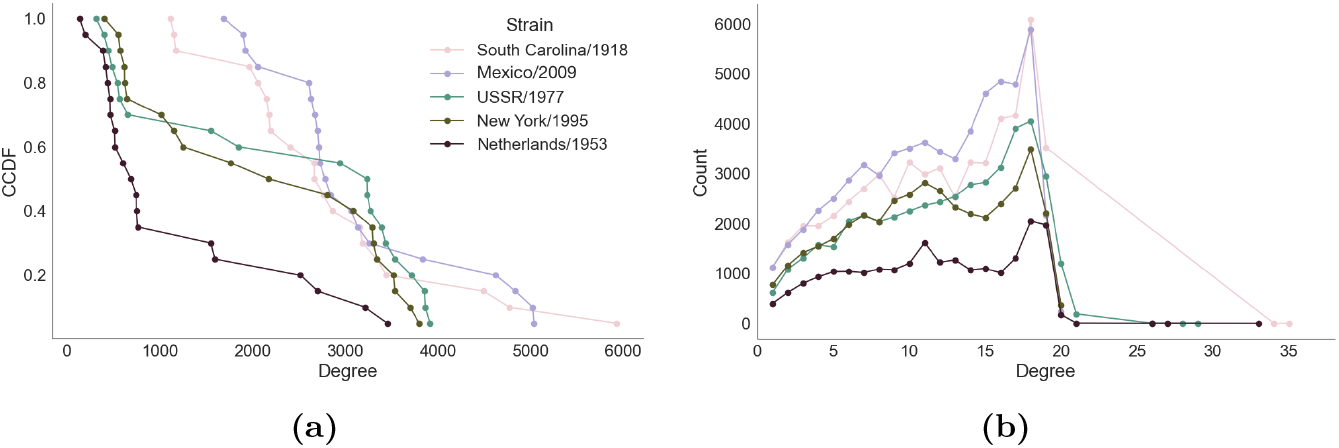
Topology of five representative NCs: (**a**) Degree distribution of seed genotypes shown as a complementary cumulative distribution function (CCDF), *P* (≥ *k*), where *k* is node degree. (**b**) Degree distribution of neighbour genotypes. Seed neighbourhoods are exhaustively sampled, whereas neighbour nodes have incomplete degree information because their own mutational neighbourhoods were not explicitly explored.

Degree distributions among neighbour nodes are comparatively limited and dominated by dense local clusters (Figure 4b). Most neighbour nodes have degrees between 10 and 20, with a peak around 18, consistent with cliques formed by positional tolerance: when a site accommodates many amino acids without structural change, all variants at that position are mutually connected. Very few neighbour nodes exceed degree 20; these likely correspond to positions tolerant across multiple overlapping seed neighbourhoods. These higher-degree nodes bridge otherwise weakly overlapping clusters, providing rare mutational paths through sequence space that would otherwise be inaccessible.

The local architecture of the HA GP map is therefore strongly position-centric. Structural neutrality is governed primarily by site-specific tolerance rather than residue identity, producing modular regions of high local connectivity. Although larger NCs exhibit higher overall genotype connectivity, traversal across the NC relies on a limited number of highly connected genotypes that bridge regions of differing positional tolerance.

### 3.3 Site-specific robustness and structural context

To investigate how mutational tolerance to structural perturbation is distributed along the HA sequence, we plotted site-specific robustness against residue position. For the full set of 197 strains, robustness values were derived from the first site-scanning loop using Equation 4 (Figure 5a). Although some strains exhibit little overall robustness, when robustness is observed it consistently localises to the same residue positions across strains, indicating conserved positional patterns of structural tolerance.

**Figure 5.**
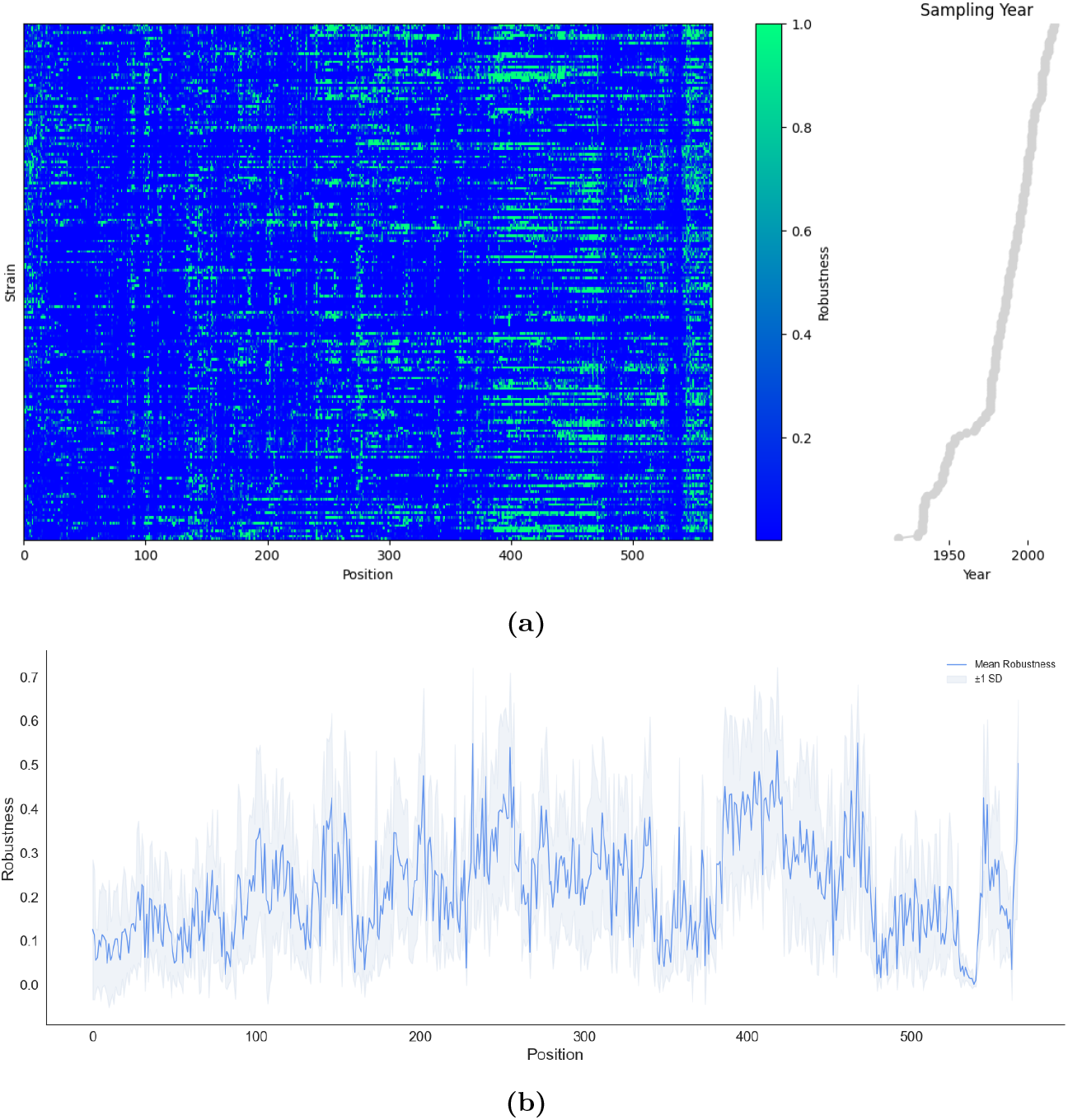
Positional robustness across HA: (**a**) Mean robustness per position across 197 strains using site-scanning, sorted by year of isolation; robustness values reflect partial amino acid sampling per site. (**b**) Mean robustness per position calculated from exhaustive single-site substitution across 15 representative strains, averaged across 20 genotypes per strain. Shaded regions denote ±1 standard deviation across strains.

Given the partial nature of site-scanning, we additionally assessed robustness using exhaustive single-site enumeration. For the subset of 15 strains, robustness was computed for each of the 20 sampled genotypes per strain using Equation 2, averaged within strains to obtain per-residue means, and subsequently averaged across strains to produce a single position-specific profile (Figure 5b). Shaded regions denote ±1 standard deviation across strains and quantify inter-strain variability in structural tolerance at each residue. Despite differences in sampling depth, the two approaches yield concordant positional profiles and reveal a strongly non-uniform distribution of robustness along the HA sequence.

This graded distribution of constraints is likely driven by functional requirements and local structural degeneracy. Regions within the stem domain (residues 385–450) exhibited high robustness, consistent with their predominantly helical structure, dense intramolecular contacts, and role in mediating post-fusion extension of the virion to the host membrane [4]. In contrast, the segment spanning residues 341–359, which includes the fusion peptide and the HA1–HA2 cleavage site, showed distinctly lower robustness, reflecting strong constraints on amino acid identity. Regions surrounding established antigenic sites [23] also displayed reduced robustness, consistent with their recurrent structural modification during immune escape.

Robustness also differed among secondary structure categories (Figure 6). Helices were significantly more robust than coils or *β*-sheets (Kruskal–Wallis test: *H* = 62.39, *p* = 2.8 × 10^−14^). This trend aligns with previous observations that helices accommodate substitutions more readily because their higher number of inter-residue contacts allows single mutations to cause less structural disruption [1].

**Figure 6.**
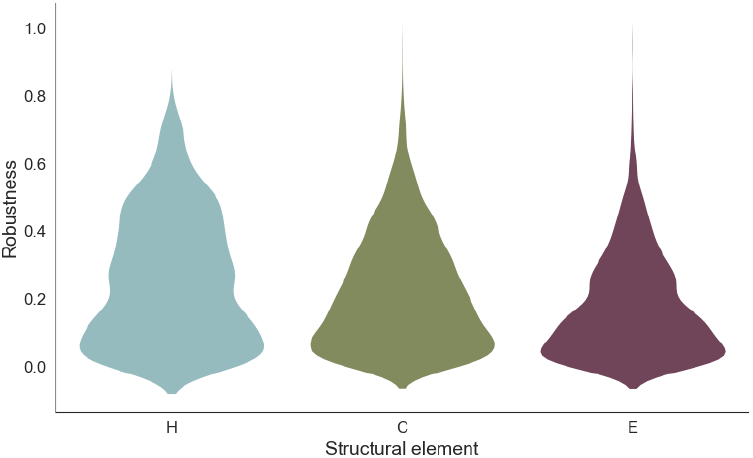
Distribution of positional robustness across secondary structure categories: Helices are significantly more robust than coils or *β*-sheets.

Together, these results indicate that robustness is modulated by both structural context and functional role. Densely packed, helical regions within the stem exhibit strong mutational buffering, whereas flexible loops and functionally critical motifs are under tighter constraint, limiting neutral exploration in those parts of sequence space.

### 3.4 Evolvability and structural transition probabilities

Evolvability is the capacity to access novel phenotypes via neutral mutations [22]. We measured it by enumerating all single–amino acid substitutions that resulted in a change of the secondary structure for the 20 genotypes sampled from each NC. This allowed us to compare the size and composition of locally accessible non-neutral phenotypes across strains.

For clarity, we define: a non-neutral neighbour genotype as a sequence that differs by a single amino acid and maps to a different phenotype; a neighbour phenotype as the phenotype of such a genotype; and a non-neighbour phenotype as any phenotype in the 2,429-structure dataset not observed among these neighbours.

In *USSR/1977*, we identified 11,363 unique neighbour phenotypes across the sampled genotypes. Among these, overlapping structures were observed in isolates from both earlier (1948–1951) and later (1978–1987) periods, with no additional overlaps detected among subsequently sampled isolates. By contrast, *Mexico/2009* exhibited 17,103 unique neighbours, indicating a larger locally accessible phenotype set. The number of overlapping structures increased in 2009, declined in 2010, and rose again in 2016. Other strains had intermediate totals (12,058–15,159).

Despite the large size of the non-neutral single-step neighbourhoods across all strains, only a minority of accessible phenotypes were unique (11–27%; Fig. 7). In all cases, the two to three most frequent non-neutral phenotypes accounted for the majority of accessible outcomes, indicating that although many structural transitions are accessible, the effective phenotypic diversity of single-step mutations remains constrained. Though strains differed subtly in the distribution of unique accessible phenotypes (Fig. 7). Mexico/2009 exhibited a higher fraction of unique accessible phenotypes, suggesting greater local evolvability relative to other strains.

**Figure 7.**
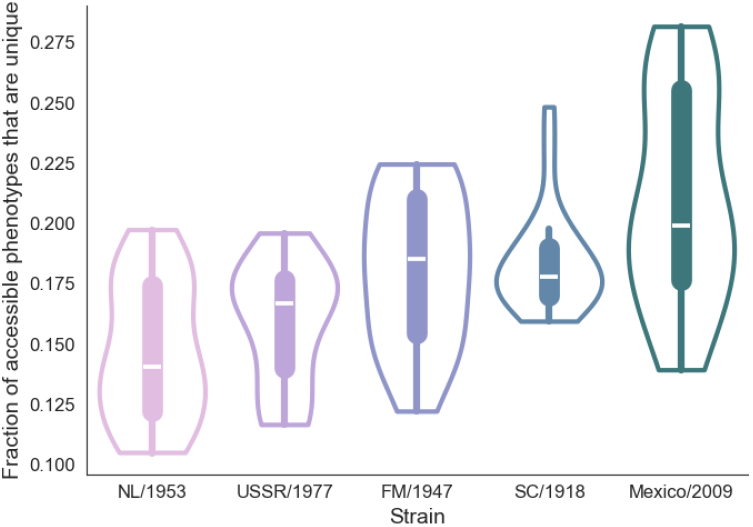
Local evolvability measured for 20 genotypes sampled from each NC: Fractions of unique and accessible phenotypes vary modestly among historical strains, with *USSR/1977* clustering around ∼ 17%. *South Carolina/1918* shows predominantly lower values with occasional higher outliers, whereas *Mexico/2009* exhibits a broader and generally elevated distribution.

To examine the relationship between NC connectivity and structural similarity, we computed the normalised Hamming distance to the reference phenotype. In *USSR/1977*, NC neighbour phenotypes were highly similar to the reference (mean distance ≈ 0.0035, s.d. ≈ 0.0018), differing at roughly 1–3 positions. Non-neighbour phenotypes were substantially more dissimilar (mean ≈ 0.028, s.d. ≈ 0.017), differing at 15 or more positions on average.

A similar pattern was observed in *Mexico/2009*. NC neighbours again showed high similarity to the reference (mean ≈ 0.003, s.d. ≈ 0.0017), while non-neighbours displayed a broader range of distances (mean ≈ 0.009, s.d. ≈ 0.012). This broader distribution reflects access to a larger set of structurally similar phenotypes and likely results in part from denser sampling of circulating strains in the post-2000 period. Since secondary structures are derived from consensus sequences in GISAID, the dataset is biased toward viruses that successfully transmitted between hosts and persisted under immune selection, while earlier periods are underrepresented, limiting neighbour and non-neighbour data for historical strains.

Although such sampling effects may inflate the tail of the non-neighbour distance distribution in recent NCs, they cannot explain the pronounced separation between neighbour and non-neighbour phenotypes. Across strains, accessible novelty is structurally proximal and heavily redundant. Together, these results indicate that evolvability in HA is a local property of the GP map, shaped jointly by NC robustness and the structural proximity of phenotypes.

## 4 Discussion and conclusion

This study provides an empirical characterisation of a protein GP map and clarifies how biophysical constraints shape evolutionary accessibility in high-dimensional amino acid sequence space. Using HA as a model system, we show that protein sequence space is partitioned into extremely uneven NCs that are locally dense yet globally sparse, and whose accessibility is strongly structured by positional tolerance. In HA, evolution therefore unfolds on a landscape dominated by strong phenotypic bias, sparse long-range connectivity, and sharply constrained routes to novelty.

This strong phenotypic bias is qualitatively similar to that observed in RNA secondary structure maps. However, unlike RNA, where large NCs typically percolate sequence space, HA NCs are embedded in an astronomically large genotype space and therefore remain boundary-dominated even at their largest observed sizes. This geometric constraint provides a parsimonious explanation for the markedly weaker scaling between robustness and NC size in proteins, with even highly robust HA structures tolerating only a small fraction of single-point mutations.

Local network reconstruction reveals that HA NCs are organised around position-centric clusters. Sites that tolerate multiple amino acids form dense local cliques, producing modular pockets of neutrality. However, overlap between these pockets is limited, and traversal across the NC appears to rely on relatively few highly connected genotypes. This architecture implies that neutral exploration in proteins is inherently heterogeneous and path-dependent. Early substitutions may irreversibly redirect accessible regions of sequence space, providing a mechanistic basis for historical contingency in protein evolution.

Structural context further modulates neutrality. Helical regions within the stem domain exhibit elevated robustness, whereas flexible loops and functionally critical motifs are more constrained. These gradients of tolerance are not captured by sequence conservation alone and instead reflect how local structural organisation buffers or amplifies mutational effects. Neutrality in proteins therefore emerges from distributed, positional tolerance rather than the compensatory base-pairing interactions of RNA folding.

Non-neutral single-step mutations are strongly biased toward structurally similar phenotypes. Most transitions converge on a small subset of outcomes, and distinct structures are rare. Evolvability is therefore predominantly incremental and redundant. Larger NCs increase the number of accessible phenotypes, but they do not alter this bias toward structurally proximate outcomes.

Together, these findings refine the scope of neutral network theory across biological systems. Core qualitative properties, phenotypic bias, robustness–evolvability coupling, and mutational accessibility, extend from RNA to proteins. However, the mechanisms generating these patterns differ fundamentally. RNA neutrality is modular and compensatory, enabling globally connected networks that facilitate long-range exploration. Protein neutrality is diffuse, site-centric, and boundary-limited. Proteins thus occupy a regime in which neutrality exists without evidence of global percolation, imposing stricter constraints on structural evolvability.

In the case of HA, these intrinsic GP map properties help explain how extensive amino acid variation can accumulate without corresponding structural divergence, and why structural innovation remains rare despite persistent immune selection. Evolutionary trajectories are shaped not only by external selective pressures but by deep geometric and biophysical constraints embedded in the protein sequence–structure map.

Several limitations should be noted. Secondary structure provides a coarse-grained phenotype, and finite local sampling necessarily underestimates long-range connectivity. Nevertheless, the consistency of patterns across strains suggests that the main conclusions are robust. Extending this framework to other proteins, incorporating higher-order structural representations, explicitly mapping multi-step neutral pathways, and integrating physics-based folding models will further clarify how protein GP maps constrain evolutionary innovation. Overall, protein sequence space supports neutrality, but in a form that is locally structured and globally constrained. Compared with RNA, this pattern suggests that neutrality in proteins may be more limited by local constraints than previously appreciated.

## Supporting information

Electronic Supplementary File

## Notes

### Competing Interest Statement

The authors have declared no competing interest.

