## Supplementary material for "Mapping protein neutral networks from predicted secondary structure": Electronic Supplementary File

### Electronic supplementary material for Mapping protein neutral networks from predicted secondary structure

#### 1 Supplementary Methods

##### 1.1 Secondary structure prediction

Predictions were performed with `Porter5` [S1, S2], which employs an ensemble of Bidirectional Recurrent Neural Networks (BRNNs) interleaved with convolutional layers to combine long-range context modelling with local motif detection. Porter5 uses MSA-derived features obtained from PSI-BLAST (iterative profile search) and HHblits (HMM–HMM comparison). Each search is run for three iterations with an $E$-value cutoff of $10^{-3}$. This cutoff retains only matches with an estimated probability ¡0.1% of occurring by chance. PSI-BLAST searches used UniRef90 and HHblits used UniProt20. Profiles from the two searches are combined to provide richer inputs than either source alone.

For each sequence, Porter5 returned a probability distribution over C, H, and E at each residue. The state with the highest posterior probability was assigned as the predicted secondary structure. Porter5 achieves approximately 84% Q3 accuracy overall (84.65% on X-ray structures and 81.6% on NMR structures) [S1], making it well suited for large-scale secondary structure phenotyping. Predicted secondary structure strings were used as phenotypic states in the GP map. Although Porter5 relies on evolutionary information, its performance is robust for HA due to the extensive representation of influenza sequences in public databases.

Predicted HA secondary structures were benchmarked against experimentally determined X-ray structures for three representative H1N1 strains with high-resolution HA structures available in the Protein Data Bank (PDB): A/California/04/2009 (3LZG) [S3], A/Fort Monmouth/1/1947 (7JPD) [S4], and A/South Carolina/1/1918 (4GXX) [S5]. Predicted sequences were trimmed according to residue-level alignment mappings obtained by structurally aligning AlphaFold2 HA models to experimental PDB structures using PyMOL's `MatchAlign` algorithm. Experimental secondary structure assignments were derived using DSSP. Predicted structures showed high concordance with experimental assignments (Q3 $\approx$ 0.74–0.77) across both HA1 and HA2 domains (Table S1).

Table S1: Structural validation of predicted HA secondary structure against experimental PDB structures. Q3 scores report residue-level secondary structure concordance between predicted and DSSP-derived assignments over aligned residues.

| Strain | PDB ID | HA1 | | HA2 | |
|---|---|---|---|---|---|
| | | Residues | Q3 | Residues | Q3 |
| *A/California/04/2009* | 3LZG | 32-333 | 0.748 | 348-518 | 0.766 |
| *A/Fort Monmouth/1/1947* | 7JPD | 23-337 | 0.740 | 348-504 | 0.732 |
| *A/South Carolina/1/1918* | 4GXX | 20-339 | 0.769 | 348-514 | 0.743 |

Evolutionary predictors such as Porter5 may underestimate the full range of structurally tolerated variation relative to physics-based folding approaches (e.g., Rosetta or explicit energy-based models). In addition, prediction runtimes (1–2 minutes per sequence) limited both sampling scale and the feasibility of extensive site-scanning analyses. More efficient predictors will be required for exhaustive characterisation of HA NCs.

#### 1.2 Size stabilisation

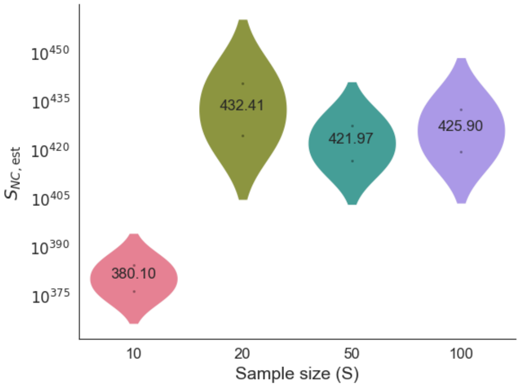


Figure S1: Stability of NC size estimates as a function of subsample size. Subsamples were drawn from a pool of approximately 1,737 putatively neutral single-point mutants for strain A/mallard/Sweden/57/2003.
